# Inferring disruption of directed graphs using LIKA reveals altered protein phosphorylation networks in schizophrenia

**DOI:** 10.64898/2026.08.06.743374

**Authors:** Lujing Zhang, Andrew G. Demarco, Kimia Ghafari, Bernie Devlin, Matthew L. MacDonald, Kathryn Roeder

**Affiliations:** Department of Statistics and Data Science, Carnegie Mellon University, 5000 Forbes Avenue, 15213, PA, United States of America; Department of Psychiatry, University of Pittsburgh School of Medicine, 3550 Terrace St, 15213, PA, United States of America; Center for Neuroscience, University of Pittsburgh School of Medicine, 3550 Terrace St, 15213, PA, United States of America; Biomedical Mass Spectrometry Center, University of Pittsburgh, 3550 Terrace St, 15213, PA, United States of America; Department of Computational Biology, Carnegie Mellon University, 5000 Forbes Avenue, 15213, PA, United States of America

**Keywords:** kinase-substrate analysis, network inference, maximum likelihood inference

## Abstract

**Motivation:** Kinases regulate a multitude of protein functions, and their dysregulation is pivotal for many human diseases. Direct measurement of kinase activity, however, is often challenging; therefore, inferring activity from the behavior of their substrates is a widely adopted strategy. Nonetheless, traditional methods typically oversimplify the underlying network, ignoring that any particular substrate can be phosphorylated by multiple kinases.

**Results:** We present *LIKA*, a likelihood-based framework for inferring kinase activity from phosphoproteomic data. By modeling the many-to-many structure of kinase-substrate interactions, LIKA achieves high efficiency, even with limited data, while capturing network complexity. Simulation and cell line analyses confirm the robustness and accuracy of LIKA. Importantly, analysis of a phosphoproteomic dataset from schizophrenia and control subjects reveals novel dysregulated kinases.

**Availability and Implementation:** The implementation code and publicly available data are provided at: https://github.com/lujingz/LIKA.

## Introduction

Protein kinases regulate a wide range of biological processes by catalyzing the addition or removal of phosphate groups on specific substrate proteins, modulating their stability, subcellular localization, enzymatic activity, and interactions with other proteins [10]. The actions of kinases have been repeatedly exploited to develop therapies against cancer, targeting individual enzymes [31]. However, because kinases operate within intricate and adaptable signaling networks to modulate cellular physiology [29], understanding the architecture and dynamics of kinase signaling networks in both health and disease are crucial for effective therapeutic strategies [29].

To this end, different approaches have been proposed to evaluate kinase networks. The most widely used approach is **KSEA** [7], which infers kinase activity by assessing the enrichment of phosphorylation levels among known substrates. It aggregates signals across substrate groups and applies statistical tests to quantify pathway activation. The method is simple and easy to implement, but it fails to account for a common case in which one or more substrates can be phosphorylated by multiple kinases. **IKAP** [21] takes into account the complexity of the network connections, but it targets the links between the kinase and phosphosites, instead of the kinases directly. Moreover, IKAP introduces a large number of parameters and imposes a higher requirement on the data, such as six or more time periods of measurement. **INKA** [5] integrates phosphosite and kinase information to improve the power of inference. However, it is limited to identifying the leading kinases in the network and cannot detect kinases that differ significantly in pairwise comparisons, which is necessary when aiming to identify disease-related kinases.

To illustrate the importance of accounting for shared substrates, we examine two networks with multiple kinases acting on a set of substrates (Fig. 1). Analyzing the phosphosites of each kinase in isolation can lead to misinterpretation: an unperturbed kinase could be falsely identified as dysregulated because it shares significant substrates with truly dysregulated kinases; while a dysregulated kinase may be overlooked if some of its substrates appear insignificant, for instance, due to the feedback regulation of a functional kinase. Ideally, a network analysis would explicitly account for the influence of other kinases, yielding more reliable and biologically meaningful results. We introduce *LIKA* (Likelihood Inference of Kinase Activity), a statistical framework for detecting dysregulated kinase activity that explicitly captures the many-to-many structure of kinase-substrate interactions. **LIKA** is data-efficient, computationally scalable, and capable of capturing the complexity of kinase-substrate networks. Through simulation studies, we demonstrate its robustness and accuracy. When applied to phospho-proteomic data from schizophrenia patients and from cell lines, **LIKA** successfully identifies disease associated dysregulated kinases in the former and dominant kinases in the latter, even under conditions of limited sample sizes and highly complex networks.

**Fig. 1.**
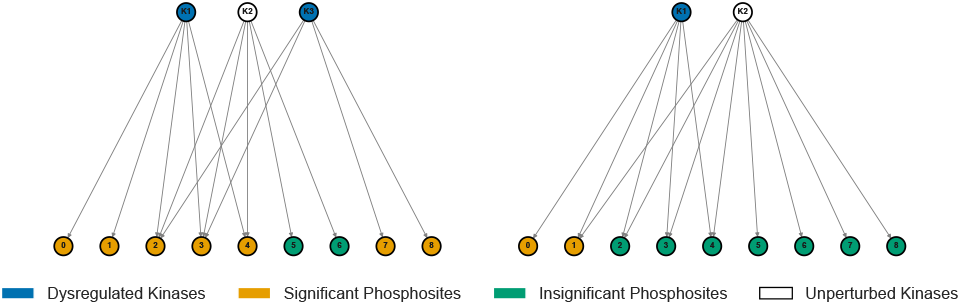
Toyexamples illustrating the limitation of ignoring the many-to many nature of kinase-substrate networks. Left: Falsely implicating an unperturbed kinase due to dysregulated phosphosites it could influence: here, many of K2’s substrates are significant solely due to the influence of K1 and K3, falsely implicating K2. Right: Overlooking dysregulated kinases for unperturbed phosphosites: many of K1 ‘s substrates are not significant because of the influence of K2, leading to missed detection of K1. Our method addresses these issues.

## Methodology

### Overview of workflow

Our procedure consists of four main steps (Fig. 2). First, we collect the raw data and process them into network data and sample data (Fig. 2, Step 1). To approximate normality, the raw intensity measurements are transformed into log-intensities. If necessary (e.g., in clinical datasets), we remove the effects of clinical covariates via regression, and use the resulting residuals as the sample data. After addressing missing values, we extract the nodes that are either measured or connected to the measured nodes from a large network database, such as KSEA [7]. Second, using the sample data, we obtain the substrate significant set (Fig. 2, Step 2); when the sample size is small, we employ the empirical Bayes shrinkage method [18] to improve statistical power. Third, combining the significant set with the network data, we estimate kinase dysregulation via the maximum likelihood method (Fig. 2, Step 3). Finally, we rank the kinases according to their estimated dysregulation levels (Fig. 2, Step 4).

**Fig. 2.**
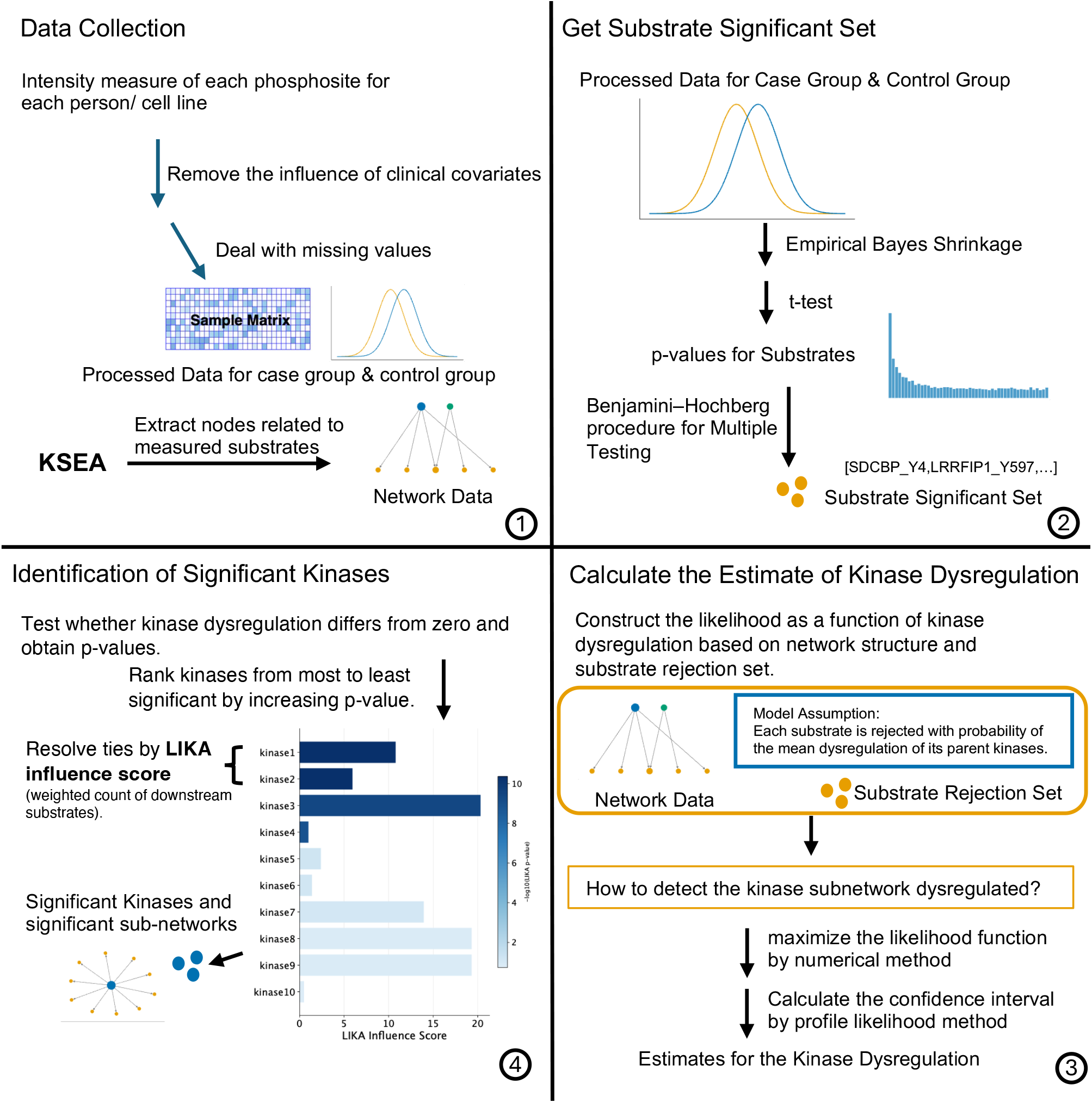
Overview of LIKA workflow. (1) Data Collection. For substrate data, we collect and preprocess the intensity measurements of each phosphosite across individuals or cell lines. For network data, we extract nodes corresponding to the measured substrates from the KSEA dataset [7]. (2) Set of significant substrates. When the number of samples is small (e.g., Experiment on cell line data), we apply empirical Bayes shrinkage to adjust the sample variance of each substrate, then compute p-values. After correcting for multiple testing using the Benjamini-Hochberg procedure, we obtain the set of significant substrates. (3) Estimation of Kinase Dysregulation. We construct a likelihood function of kinase dysregulation based on the network structure and the set of significant substrates. The likelihood is maximized numerically, and profile likelihood is used to derive confidence intervals for each kinase. (4) Identification of Significant Kinases. For each kinase, we test whether its dysregulation differs from zero and obtain a kinase-level p-value. Kinases are ranked from most to least significant by increasing p-value. Ties - and, in our recommendation, any set of kinases assigned very small p-values - are resolved using the LIKA influence score, defined as a weighted count of downstream substrates. The top-ranked kinases, together with their associated substrates, define the significant dysregulated subnetworks.

### Data Collection

The analysis requires two types of input data. First, a kinase-substrate network is used to define curated regulatory relationships derived from phosphoproteomics studies. This network is represented as a directed graph, where nodes correspond to kinases or substrates and edges indicate kinase-substrate regulatory relationships. Second, phosphosite-level measurements from two groups of samples are used to quantify differential phosphorylation activity. To reduce potential confounding, the effects of relevant clinical covariates are adjusted for prior to downstream analysis.

### Get Substrate Significant Set

Under the assumption of approximate normality, we employ per-substrate *t*-tests based on the sample mean difference and variance to obtain *p*-values. In small-sample contexts, however, as commonly encountered in cell line analyses, variance estimates for individual substrates are highly variable, which can reduce statistical reliability.

To address this issue, we employ an empirical Bayes shrinkage method [18], which borrows strength across substrates to stabilize variance estimates and thereby improve the reliability of the test statistics. Based on the resulting per-substrate *p*-values, we then apply the Benjamini-Hochberg procedure [6] to identify significant substrates, although alternative multiple testing corrections could also be considered.

### Maximum Likelihood Estimation of Kinase Dysregulation

Based on biological intuition, we assume that each substrate is significantly perturbed with a probability equal to the mean dysregulation of its parent kinases, where the dysregulation levels are unknown and must be inferred from the data. To achieve this, we formulate a likelihood function in terms of kinase dysregulation and estimate the parameters using the maximum likelihood method.

Let *K* and *S* denote the sets of kinases and substrates, respectively. For *v ∈ K*, let *8(v)* represent the dysregulation level of kinase *v*. For *s ∈ S*, let Parents(s) be the set of its parent kinases. We assume that the probability of substrate _*s*_ being significant is given by the mean dysregulation of its parent kinases:

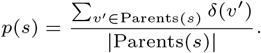

Here | Parents(*s)*| is the count of possible kinase parents or degree of the node *s*, which we will rewrite as *Ds*. Let *r(s)* be the indicator variable denoting whether the substrate _*s*_ is significant according to the significant set. The joint likelihood of the significant data {*r(s)* : *s ∈* S} is given by

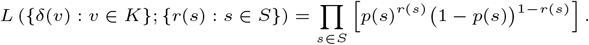

We estimate the unknown dysregulation levels {*δ* (*v)* : *v* ∈ *K* } by maximizing this joint likelihood. Because no closed-form solution exists, the optimization is performed numerically.

### Identification of Significant Kinases and Significant Subnetworks

LIKA compares each kinase’s dysregulation against the background significance rate set by the substrate-level Benjamini-Hochberg procedure. If δBH is the realized raw *p* value cutoff at FDR 0.05, then under the null a substrate is significant with probability approximately δBH; LIKA therefore uses δBH as the null dysregulation level.

For kinase *v* ∈ *K*, let *δ-v* = *{δ (u): u ∈ K*\ {v}} denote all other kinase dysregulation levels. We profile out these nuisance parameters:

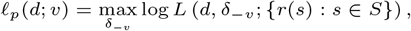

where *d* is the value assigned to *δ (v)* and *r(s)* indicates whether substrates *s* is significant. Let 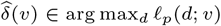. We test *Ho* : *δ(v)* = ^*δ*^BH versus *HA* : *δ (v)* > ^*δ*^BH, using the signed likelihood-ratio statistic 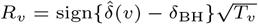.

Under standard regularity conditions, *R*_*v*_ is asymptotically standard normal under *H*_*o*_ [4]. Therefore, the one-sided kinase level p-value is *P*_*v*_ = 1 - *Φ (R*_*v*_).

Kinases are ranked by increasing kinase-level p-value. For very small p-values, we additionally prioritize kinases by downstream influence. Specifically, we use the truncated p value

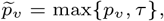

where *T* may be chosen as a Bonferroni-corrected significance threshold, and break ties using the LIKA influence score

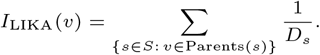

This score is the effective number of downstream substrates assigned to kinase *v*: substrates with fewer candidate parent kinases receive greater weight, while shared substrates are not over-counted. For each selected kinase, the significant subnetwork consists of the kinase, its significant downstream substrates, and the corresponding kinase-substrate edges.

## Experiments

We conducted three sets of experiments: a simulation study; phosphorylation levels detected in cell line data, and in which our analysis contrasts levels between two cell lines; and an analysis of differential mean phosphorylation levels detected in postmortem brain samples from schizophrenia subjects versus controls. The simulation study demonstrates our model’s performance on complex networks. The cell line analysis, for which ground-truth knowledge is well established, validates our model’s ability to identify dysregulated kinases in real data and, importantly, to recover downstream kinases of the dominant kinase. Finally, the schizophrenia study provides biological insights into potential mechanisms underlying the disorder. The code supporting all experiments is available at https://github.com/lujingz/LIKA.

### Simulation Results

We evaluated LIKA using simulated kinase-substrate networks with known dysregulated kinases. Substrate-level significance was generated from a binomial model in which the probability of significance depended on the dysregulation levels of the substrate’s parent kinases. Significant substrates were assigned a case-control shift, with case intensities sampled from *N*(2, 1) and control intensities from *N*(O, l); non-significant substrates were sampled from *N*(0, 1) in both groups. Each group contained five samples, creating a small-sample setting for kinase inference.

We considered three network structures: non-overlapping substrates with equal kinase degree, non-overlapping substrates with unequal kinase degree, and overlapping substrates with unequal kinase degree. Detailed network constructions are provided in Supplementary Material 1, and the simulated network structures are summarized in Supplementary Table S1. KSEA was applied to the same datasets as a baseline (Fig. 3).

**Fig. 3.**
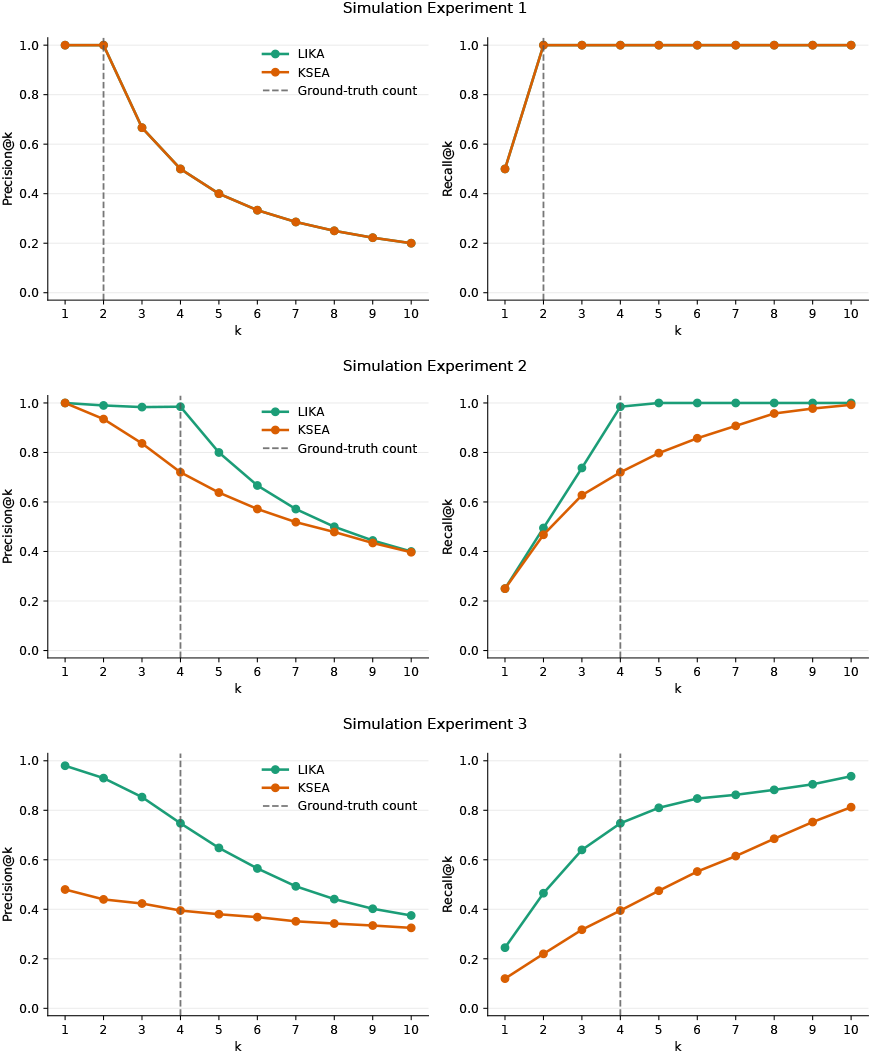
Simulation performance under three synthetic kinase-substrate network structures. Precision@k (left) and recall@k (right) are averaged over 100 runs; kinases are ranked by capped LIKA p-value or KSEA *p* value, and dashed lines indicate the true number of dysregulated kinases. LIKA matches KSEA in the balanced setting and outperforms it under degree heterogeneity and substrate overlap.

Performance was measured by precision@k and recall@k. Let *G* be the set of ground-truth dysregulated kinases and *T*_*k*_ the top *k* predicted kinases:

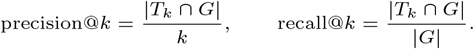

In the simplest setting, LIKA and KSEA both recovered all ground-truth kinases at the true kinase count (Fig. 3). When kinase substrate counts were unequal, LIKA maintained perfect recovery, whereas KSEA showed reduced prec1s1on and recall. Under both degree heterogeneity and substrate overlap, LIKA consistently outperformed KSEA across a broad range of *k*, demonstrating greater robustness to heterogeneous and shared kinase-substrate annotations.

### Experiment on cell line data 5

To illustrate our method when we know the ground truth, we utilize data from two cancer cell lines - K562 and HCC827-ER3 [5] - treating these cell lines as case-control samples to determine which kinases are identified by the contrast of phosphorylation levels found on proteins in the cell lines. The key mutation of K562 is a fusion of chromosomes 9 and 22, generating the fusion gene *BCR-ABL1* [27]. *BCR ABL1* substantially elevates tyrosine kinase activity, as well as altering activity of various kinases (Serine/threonine-protein kinase CRK1, phosphoinositide 3-kinase PI3K) and pathways (RAS/RAF /MAPK, PI3K/ AKT/mTOR, JAK/STAT, and WNT/ β-catenin) [2]. The HCC827-ER3 cell line carries a heterozygous in-frame deletion of 5 amino acids (p.Glu746-Ala750del) in *EGFR*, a receptor tyrosine kinase, and also carries a homozygous in-frame mutation (p.Val218del) in *TP53*. The mutation in *EGFR* activates the Ras/mitogen-activated protein kinase and PI3K/ Akt pathways [13]. Although *BCR ABL1* and *EGFR* overlap in some activation pathways, these cell lines should differ substantially in the phosphorylation patterns of their substrates.

An important feature of this experiment is the sample size. Although most case-control studies have relatively large sample sizes, the data produced by Beekhof et al. [5] are limited to phosphorylation measures from two samples of each cell line, K562 and HCC827-ER3, and a technical replicate for each sample. However, because the variability of measurements over samples of the same cell line is low, we expected the data to contain sufficient signal. The two cell lines share 504 phosphosites, 296 of which are recorded in KSEA [7]. We therefore limited our analyses to kinases associated with these 296 phosphosites, from which we constructed a network comprising 365 nodes. Each phosphosite has four measurements per cell line, with partial missingness. The log-transformed intensities, which are approximately normally distributed, were used as input data. An empirical Bayes method [18] was used to contrast phosphosite values. Additional preprocessing details and substrate-level results are provided in Supplementary Material 2 and Supplementary Table S3. Both EGFR and ABL1 are among the top 10 kinases detected by LIKA score (Fig. 4), while only ABL1 is detected by KSEA, highlighting the sensitivity of LIKA. In addition, beyond the two focal kinases, LIKA identifies 4 kinases that are direct downstream targets of these focal kinases, namely FYN, SRC, LYN, and HCK, whereas KSEA identifies none. Notably, both kinases regulate a large number of substrates (Fig. 5), most of which exhibit significant differences between the two cell lines. The substrates with the smallest p-values are labeled, forming an “extremely hot” region.

**Fig. 4.**
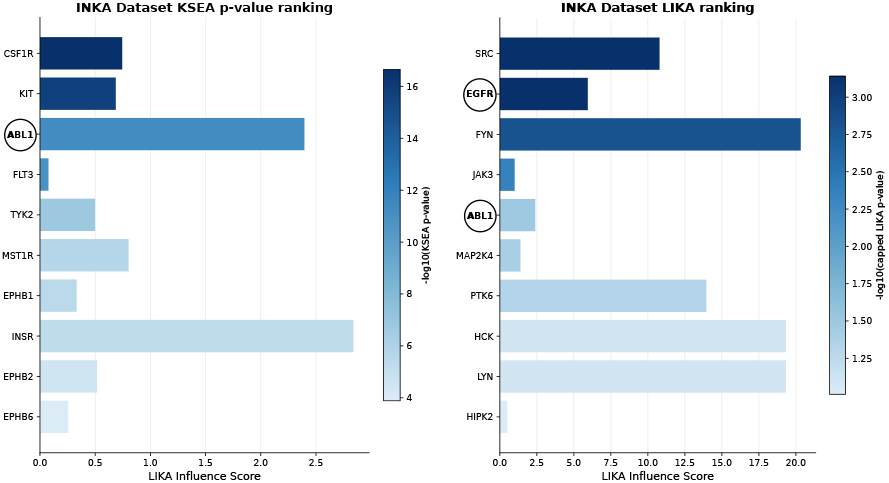
Results for the INKA dataset. Left: Top ten kinases ranked by KSEA p-value. Right: Top ten kinases ranked by LIKA p-value. Kinases are ordered from rank 1 to rank 10 from top to bottom. Bar length shows the LIKA influence score, and color intensity shows −log_10_(*p*-value), with darker colors corresponding to smaller p-values. Ground-truth kinases reported by Beekhof et al. [5] are circled on the y-axis. LIKA identifies both EGFR and ABLl among the top ten kinases, whereas KSEA identifies only ABLl, suggesting that LIKA better recovers the reported ground truth in this example. Complete kinase rankings and associated statistics are provided in Supplementary Table S2.

**Fig. 5.**
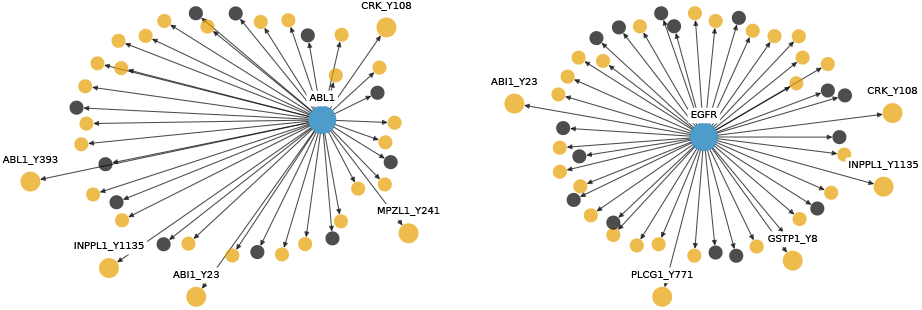
Subnetworks associated with ABL1 and EGFR. Yellow nodes represent significant substrates, whereas black nodes represent insignificant substrates. Among the yellow nodes, those with labels correspond to the five most strongly significant substrates, characterized by the smallest p-values.

### Perturbation of kinase function in schizophrenia

#### Dataset Overview

For this experiment, a postmortem sample from the dorsal anterior cingulate cortex (dACC) of each of 112 subjects was assessed by TMT-labeled mass spectrometry for phospho-proteomic detection and measurement. Of the 112, 56 of the subjects were diagnosed with schizophrenia. These subjects were pair-matched to 56 subjects who had no psychiatric diagnosis, matching on sex, age band, and postmortem interval (PMI). Pairs were then processed together for all experimental protocols. Of the 45,471 phosphopeptides detected as greater than zero abundance, the abundances were confidently estimated for 18,372. To estimate the mean phosphorylation differences between cases and controls, we first subjected these estimates to a model selection procedure in which covariates were retained if they were useful predictors for at least 5% of the phosphopeptides. The final model included case-status, as well as covariates plex (TMT batch), PMI, age, ancestry, and res.pH (residual obtained after regressing pH on diagnosis). However, of these phosphopeptides, 1,740 could be confidently assigned to a specific phosphosite that was also recorded in KSEA (Supplementary Table S4), and these were analyzed here. Of these phosphosites, 1,231 had significantly different means (FDR < 0.05) for cases versus controls.

Next, based on the 212 kinases associated with these 1,740 phosphosites, we constructed a network comprising 1,952 nodes (Supplementary Table S4). When kinases share identical substrates, it is impossible to determine which one drives the effect. To address this challenge, we merged such kinases by treating them as a single node in the network, with the union of their original parents and children assigned as the parents and children of the merged node.

#### Main findings

Using the kinase network and the p-values from the case control contrast, LIKA and KSEA highlighted many of the same kinases as prominently differentiated (Fig. 6, Supplementary Table S5). Of the top 10 for each method, four overlapped, namely MAPK3, CSNK2Al, CDKl, and CDK2. Rare mutations in *CSNK2A1* have been weakly associated with schizophrenia ([25]), although it is strongly associated with other neurodevelopmental disorders ([25]). The different kinases in the top 10 sets were revealing about the methods. For example, while KSEA highlighted NEK3, it had only one very significant substrate, `SUGT1_T265. SUGT1_T265` was also a target for 8 other kinases, including CDK1, CDK2 and MAPK3, which were also in the KSEA top 10 set. Thus, it is plausible that the differential case-control phosphorylation signal at `SUGT1_T265` arose from the action of other top kinases, rather than NEK3. Another observation of interest: CDK1 and CDK2 shared over half of their significant substrates with MAPK3. In fact, MAPK3 (also known as ERK1) was a plausible actor for much if not all of the relevant differential phosphorylation targets, in the sense that non-coding variation impacting expression of this kinase has been implicated by early (Supplementary Information and Data file 2, [11]) and recent (Supplementary Table 12 of [30]; see also [9]) colocalization of genome-wide association (GWAS) findings for schizophrenia. *MAPK3* falls in chromosome region 16p11.2, which is subject to copy number variation: duplications of this region elevate risk for schizophrenia ([20]), whereas deletions and sometimes duplications confer risk for autism spectrum disorder [15, 32, 19]. Notably, the GWAS-discovered common risk variation also elevates expression of *MAPK3*.

**Fig. 6.**
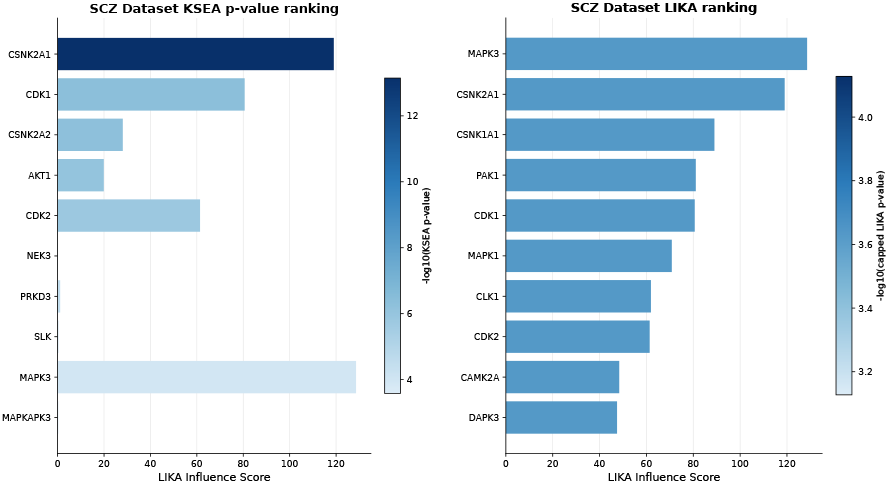
Results for the schizophrenia dataset. Left: Top ten kinases ranked by KSEA p-value. Right, Top ten kinases ranked by LIKA p value. In each panel, kinases are ordered from rank 1 to rank 10 from top to bottom. The bar length represents the LIKA influence score, while the color intensity represents the corresponding −log_10_(p-value), with darker colors indicating smaller p-values. When multiple kinases have the same p value, ties are broken by the LIKA influence score, with the kinase having the larger score ranked higher. Complete kinase rankings and associated statistics are provided in Supplementary Table S5.

Another interesting contrast of the two methods arises for TTBK2, which showed strong association by LIKA, and SLK, which ranked highly by KSEA. Both kinases had three substrates showing differential phosphorylation. The substrates putatively phosphorylated by TTBK2, however, had no other known kinases that could phosphorylate the sites. By contrast, the substrates putatively phosphorylated by SLK could be phosphorylated by many other kinases. For this reason, LIKA reported a p-value of association of 0.40, whereas KSEA reported a highly significant p-value of association, 5.56 × 10^−5^ A sensitivity analysis of the first-stage FDR threshold is provided in Supplementary Material 3.

## Conclusion

Motivated by the limitations of existing kinase inference approaches, we developed **LIKA**, a likelihood-based framework that enables robust inference of kinase activity and dysregulation, accounting for the complexity of phospho-signaling networks. By explicitly modeling the many-to-many structure of kinase substrate relationships, **LIKA** overcomes key limitations of simple enrichment-based approaches and provides a more realistic representation of biological signaling. Our results across simulation,cell line,and schizophrenia datasets demonstrate that incorporating network structure into statistical inference substantially improves both the accuracy and interpretability of kinase activity estimates.

One of the main strengths of LIKA is its ability to disentangle overlapping kinase-substrate relationships, a pervasive feature of cellular signaling systems. Protein phosphorylation networks are inherently redundant and highly interconnected with individual phosphosites often regulated by multiple kinases depending on cellular environment [16]. Traditional approaches such as KSEA implicitly assume independence among substrates, which can lead to systematic biases when substrates are shared across kinases. By contrast, LIKA formulates kinase inference as a global optimization problem, which allows to partition shared substrate signals across multiple candidate kinases. This results in improved robustness in complex network settings, as demonstrated in our simulation studies.

Our application to cell line data highlights LIKA’s ability to identify not only primary driver kinases but also downstream signaling components. In the comparison between K562 and HCC827 cell lines, LIKA identified both the oncogenic drivers (BCR-ABL1 and EGFR), along with several Src family kinases (FYN, SRC, LYN, and HCK). This observation is consistent with extensive literature demonstrating that oncogenic kinases propagate signals through downstream kinase cascades, particularly via Src-family kinases, which serve as central hubs in tyrosine kinase signaling [23, 33]. This successful identification of well documented downstream effectors indicates that LIKA captures not only first-mover kinase but also higher-order network propagation effects, critical for understanding pathway-level dysregulations.

In our analyses of the schizophrenia versus control differential phosphorylation, both LIKA and KSEA converge on four kinases in their top 10, namely MAPK3, CSNK2Al, CDKl, and CDK2. P-values for the LIKA top ten are uniformly small, and therefore we rank kinases by the LIKA influence score. For KSEA, p-values for these kinases are also small, the largest being MAPKlO at p-value = 0.0024. Nonetheless, the rankings with these kinases can be very different; LIKA places MAPK3 first by LIKA influence score (Fig. 6); whereas KSEA ranks it in the ninth position by p-value (Supplementary Table S5). In either case, MAPK3 is of keen interest because genetic studies have implicated increased MAPK3 expression in schizophrenia risk, including evidence from 16.pll.2 duplication studies and transcriptomic analyses prioritizing MAPK3 as a candidate susceptibility gene [11, 30, 20]. For this reason, we asked whether potential phosphosites targeted by MAPK3 show greater levels of phosphorylation in SCZ than controls, on average? The distribution (Supplementary Fig. SlA) of differential phosphorylation MAPK3 phosphosites shows a long right tail, consistent with genetic evidence implicating MAPK3-associated signaling in schizophrenia. Nonetheless, and intriguingly, both tails are heavier than expected, suggesting MAPK3-associated signaling in brain of individuals with schizophrenia is more complex than what would be predicted from the genetic association alone and may reflect substrate-specific or context-dependent remodeling of ERK signaling.

While its association with schizophrenia is not established with certainty [25, 28], CSNK2A1 has profound impact on neurodevelopment. Mutation of one copy of the gene can give rise to intellectual disability, behavioral problems, speech problems, and microcephaly, among other developmental challenges [22]. Such mutations are also associated with autism spectrum disorder [12], which is genetically correlated with schizophrenia [3, 26, 14]. CSNK2A1 is a component of the catalytic domain of casein kinase 2 (CK2), as is CSNK2A2. CSNK2A2 is scored highly by LIKA and KSEA: LIKA yielding a p-value of 1.1 × 10^−16^; KSEA yielding a p-value of 5.9 × 10^−7^. In fact, CK2 is a tetramer consisting of two active subunits, either homo- or heterodimers of these two proteins, with CSNK2B serving as the two regulatory subunits [17]. That both active domain subunits are implicated in schizophrenia by LIKA and KSEA is therefore intriguing: perhaps CK2, when composed of the heterodimer active domain, plays an important regulatory role affecting development of schizophrenia. Contrary to MAPK3, the phosphosites for both CSNK2A1 and CSNK2A2 tend to be uniformly dephosphorylated in subjects with schizophrenia (Supplementary Fig. S1B-C). Postmortem studies have reported reduced CK2 levels and activity in cortex in subjects with schizophrenia, including reduced phosphorylation of the CK2 substrate syntaxin-1 [8]; [1], making the tendency toward lower phosphorylation of predicted CK2 substrates remarkably consistent with prior observations.

Notably, CK2 and ERK1 are mechanistically linked. CK2 regulates the nuclear sstranslocation and activity of ERK1 and ERK2 through phosphorylation of key serine residues required for importin-7 binding [24]. ERK2 (MAPK1) was also among the top kinases identified by LIKA. The simultaneous implication of MAPK3, MAPK1, and both CK2 catalytic subunits therefore raises the possibility that these findings reflect perturbations within an interconnected signaling network rather than independent kinase abnormalities.

From a methodological standpoint, the stability of LIKA across a wide range of hyperparameter choices further underscores its robustness. The observed consistency in kinase rankings across varying FDR thresholds and confidence interval lengths suggests that the inferred signaling patterns reflect underlying biological structure rather than artifacts of statistical tuning. This robustness is particularly important in high-dimensional phosphoproteomics data, where arbitrary thresholding can substantially impact downstream interpretation.

Overall, LIKA provides a statistically rigorous and biologically informed framework for kinase inference in complex signaling networks. By integrating network topology with probabilistic modeling, it enables more accurate identification of dysregulated kinases and their associated pathways. Its application to schizophrenia data links genetic variation affecting MAPK3 (ERK1) to its impact on differential phosphorylation in brain tissue, offering a more integrated view of disease-related signaling alterations. More broadly, LIKA represents a generalizable approach for dissecting signaling network perturbations across diverse biological contexts.

## Competing interests

No competing interest is declared.

## Data and Code Availability

The source code, documentation, and scripts required to reproduce the analyses are available at https://github.com/lujingz/LIKA. The repository includes the implementation of LIKA, scripts for the simulation studies, and scripts for reproducing the cell-line and schizophrenia analyses described in this article.

The kinase-substrate network used in this study was obtained from the KSEA web resource https://casecpb.shinyapps.io/ksea/. The cell-line phosphoproteomic data were obtained from Supplementary Dataset EV2 of Beekhof et al. [5]. Processed substrate-level statistics, kinase substrate annotations, LIKA rankings, KSEA rankings, and supplementary results are provided in Supplementary Tables S2-S5.

The processed phosphosite-level input data used for the LIKA schizophrenia analysis, including phosphosite identifiers, case-control summary statistics, substrate-level p-values, significance indicators, and upstream kinase annotations, are provided in Supplementary Table S4 and in the accompanying code repository. These processed data are sufficient to reproduce the reported kinase-level LIKA analyses. Access to additional subject-level or full phosphoproteomic data may be requested from the corresponding authors and will be considered subject to institutional approval and applicable data-use restrictions.

## Author contributions statement

Lujing Zhang (Conceptualization [equal], Formal analysis [lead], Methodology [equal], Software [lead], Validation [equal], Visualization [lead], Writing—original draft [equal], Writing — review and editing [equal]); Andrew G. Demarco (Data curation [equal], Validation [supporting], Writing —original draft [supporting], Writing — review and editing [supporting]); Kimia Ghafari (Data curation [equal], Validation [supporting], Writing—original draft [supporting], Writing—review and editing [supporting]); Bernie Devlin (Conceptualization [equal], Supervision [equal], Validation [equal], Writing—original draft [equal], Writing—review and editing [equal]); Matthew L. MacDonald (Conceptualization [equal], Resources [lead], Supervision [equal], Investigation [equal], Validation [equal], Writing—review and editing [equal]); Kathryn Roeder (Conceptualization [equal], Methodology [equal], Supervision [equal], Validation [equal], Writing—original draft [equal], Writing—review and editing [equal]).

## Acknowledgments

This work is supported in part by funds from the Simons Foundation (SF1018804) and the National Institutes of Health (NIMH: MH125235, MH123184).

## Notes

https://casecpb.shinyapps.io/ksea/

https://pmc.ncbi.nlm.nih.gov/articles/PMC6461034/#_ad93_

### Competing Interest Statement

The authors have declared no competing interest.

