## Supplementary Material for "Inferring disruption of directed graphs using LIKA reveals altered protein phosphorylation networks in schizophrenia"

### Supplementary Material 1: Simulation Details

This section provides additional details on the simulated kinase–substrate networks used to evaluate the performance of the proposed method. The simulations were designed to assess whether the method can recover dysregulated kinases under different network topologies, including settings with balanced kinase degrees, heterogeneous kinase degrees, and overlapping substrate annotations. Each simulated dataset contains two groups: a control group and a case group. For every substrate, five control intensities and five case intensities were generated. Control samples were always drawn from a standard normal distribution, whereas case samples were shifted only for substrates selected as perturbed by the dysregulated kinases.

Kinase–substrate networks were represented as directed bipartite graphs. Each edge was directed from a kinase to a substrate, so an edge $K_i \to s_j$ indicates that kinase $K_i$ regulates substrate $s_j$. The input network table therefore contains one row per kinase–substrate edge, with a kinase identifier in the `from` column and a substrate identifier in the `to` column. Kinases were named $K0, K1, \ldots$, and substrates were named by integer identifiers. In the first two simulations, substrate sets were non-overlapping, so every substrate had exactly one upstream kinase. In the third simulation, substrate sets were allowed to overlap, so a substrate could have multiple upstream kinases.

We considered three simulated network structures:

- **Simulation 1: non-overlapping network with equal kinase degree.** This network contains 10 kinases, named $K0$ to $K9$. Each kinase is connected to 10 distinct substrates, resulting in 100 substrates in total. The substrate sets are non-overlapping: $K0$ regulates substrates 0–9, $K1$ regulates substrates 10–19, and so on. Thus, each substrate is regulated by exactly one kinase. We set $K0$ and $K1$ as the dysregulated kinases. Accordingly, substrates 0–19 were perturbed in the case group, whereas substrates 20–99 were not perturbed.
- **Simulation 2: non-overlapping network with unequal kinase degree.** This network contains 13 kinases, named $K0$ to $K12$, with heterogeneous numbers of downstream substrates. The kinase substrate counts are

  $$20,\ 20,\ 15,\ 15,\ 12,\ 11,\ 10,\ 8,\ 6,\ 4,\ 3,\ 2,\ 1,$$

  respectively, resulting in 127 distinct substrates in total. Substrates are assigned consecutively to kinases without overlap. For example, $K0$ is connected to the first 20 substrates, $K1$ to the next 20 substrates, and so forth. We set $K0$, $K2$, $K5$, and $K8$ as the dysregulated kinases. Because every substrate has exactly one upstream kinase in this simulation, all substrates downstream of these dysregulated kinases were perturbed, while substrates downstream only of non-dysregulated kinases were not perturbed.
- **Simulation 3: overlapping network with unequal kinase degree.** This network also contains 13 kinases, named $K0$ to $K12$, with the same kinase substrate counts used in Simulation 2:

  $$20,\ 20,\ 15,\ 15,\ 12,\ 11,\ 10,\ 8,\ 6,\ 4,\ 3,\ 2,\ 1.$$

  In contrast to the first two simulations, all kinase substrate sets were sampled from a shared pool of 40 substrates. For each kinase, the specified number of substrates was sampled without replacement from this pool. Because different kinases sampled from the same substrate pool, the same substrate could be regulated by multiple kinases. This setting represents the many-to-many regulatory relationships commonly observed in kinase–substrate networks. We again set $K0$, $K2$, $K5$, and $K8$ as the dysregulated kinases.

#### Generation of perturbed substrates

After constructing each network, we determined which substrates were perturbed in the case group. Let $\mathcal{P}(s)$ denote the set of upstream kinases for substrate $s$, and let $\mathcal{D}$ denote the set of dysregulated kinases. For Simulations 2 and 3, the probability that substrate $s$ was perturbed was defined as

$$\Pr(s \text{ is perturbed}) = \frac{|\mathcal{P}(s) \cap \mathcal{D}|}{|\mathcal{P}(s)|}.$$

This rule has a simple interpretation. If all upstream kinases of a substrate are dysregulated, the substrate is always perturbed. If none of its upstream kinases are dysregulated, the substrate is never perturbed. If a substrate is shared by both dysregulated and non-dysregulated kinases, the perturbation probability is proportional to the fraction of its upstream kinases that are dysregulated. This construction aligns with the modeling assumption used by LIKA: the observed significance status of a substrate is treated as arising from the combined influence of its upstream kinases. In particular, when a substrate has multiple candidate upstream kinases, its perturbation probability depends on how many of those upstream kinases are truly dysregulated, rather than assigning the substrate exclusively to a single kinase. Therefore, the simulation directly tests whether LIKA can use shared substrate information to recover the underlying dysregulated kinases in a setting consistent with its many-to-one kinase–substrate likelihood model.

In Simulation 2, every substrate has exactly one upstream kinase, so this rule reduces to a deterministic assignment: substrates downstream of $K0$, $K2$, $K5$, or $K8$ are perturbed with probability 1, and all other substrates are perturbed with probability 0. In Simulation 3, substrates may have multiple upstream kinases, so perturbation assignment is stochastic for substrates shared by both dysregulated and non-dysregulated kinases. This design makes Simulation 3 more challenging because a perturbed substrate may be connected to several candidate kinases, not only to the true dysregulated kinase.

Simulation 1 used a deterministic perturbation design. Since $K0$ and $K1$ were specified as dysregulated and each has 10 non-overlapping substrates, substrates 0–19 were perturbed and substrates 20–99 were unperturbed.

### Generation of case and control intensity data

For each substrate, we generated two rows in the simulated intensity matrix: one row for the control group and one row for the case group. Each row contains five replicate intensity measurements. The control group was generated as

$$X^{(0)}_{s,r} \sim \mathcal{N}(0,1), \qquad r = 1, \dots, 5,$$

for every substrate $s$. The case group was generated according to whether the substrate was selected as perturbed:

$$X^{(1)}_{s,r} \sim \begin{cases} \mathcal{N}(2,1), & \text{if substrate } s \text{ is perturbed,} \\ \mathcal{N}(0,1), & \text{otherwise.} \end{cases}$$

Thus, perturbed substrates have an expected case-control shift of 2, whereas unperturbed substrates have no expected difference between the two groups. No log transformation was applied to the simulated intensities when running LIKA or KSEA.

### Repeated simulation runs and evaluation

For the repeated simulation evaluation, each experiment was run 100 times using different random seeds. The random seed controls the generated control and case intensities in all simulations. In Simulations 2 and 3, it also controls the stochastic selection of perturbed substrates. In Simulation 3, it additionally controls the random assignment of substrates to kinases, because the overlapping network is generated by sampling substrate sets from a shared pool.

For each run, both LIKA and KSEA were applied to the same simulated network and intensity matrix. LIKA candidates were ranked by kinase-level LIKA $p$-value, with ties broken by the number of efficient substrates. KSEA candidates were ranked by increasing KSEA $p$-value. For each method, the top $k$ predicted kinases were compared against the known dysregulated kinase set. Let $P_k$ denote the set of top $k$ predicted kinases and let $D$ denote the set of true dysregulated kinases. Precision@k was defined as

$$\text{Precision@}k = \frac{|P_k \cap D|}{|P_k|}.$$

This quantity measures the fraction of selected candidates that are true dysregulated kinases. A high precision@k indicates that the top-ranked list has few false positives. Recall@k was defined as

$$\text{Recall@}k = \frac{|P_k \cap D|}{|D|}.$$

This quantity measures the fraction of all true dysregulated kinases recovered among the top $k$ predictions. A high recall@k indicates that the method places most or all true dysregulated kinases near the top of the ranking. Precision and recall emphasize complementary aspects of ranking quality: precision is most informative about the reliability of a short candidate list, whereas recall is most informative about whether the full ground-truth signal has been captured by a given cutoff. We calculated both metrics for $k = 1, \dots, 10$ and averaged them across the 100 simulation runs.

Table S1 summarizes the number of kinases, the number of distinct substrates, the dysregulated kinases, and whether substrate overlap is present in each simulated network.

| Simulation setting | Number of kinases | Number of distinct substrates | Substrate overlap | Dysregulated kinases |
|---|---|---|---|---|
| #1, non-overlapping, equal-size | 10 | 100 | No | $K0, K1$ |
| #2, non-overlapping, unequal-size | 13 | 127 | No | $K0, K2, K5, K8$ |
| #3, overlapping, unequal-size | 13 | 40 | Yes | $K0, K2, K5, K8$ |

**Table S1.** Summary of the simulated kinase–substrate network structures. Simulation 1 evaluates performance under a balanced network topology in which all kinases have the same number of non-overlapping substrates. Simulation 2 introduces heterogeneity in the number of substrates regulated by each kinase while preserving non-overlapping substrate sets. Simulation 3 uses the same heterogeneous kinase degrees as Simulation 2 but samples downstream substrates from a shared substrate pool, allowing substrates to be regulated by multiple kinases.

### Supplementary Material 2: Cell-line Experiment Details

For the substrate-intensity data, we downloaded Supplementary Dataset EV2 from Beekhof et al. [35] For kinase–substrate relationships, we used the network distributed with the KSEA method, downloaded from https://casecpb.shinyapps.io/ksea/. We excluded phosphosites that could not be mapped to the KSEA network and phosphosites that were not observed in either of the two cell lines analyzed here, K562 and HCC827-ER3.

Because no more than four samples were available for each cell line, substrate-specific variance estimates from ordinary $t$-tests are expected to be noisy and sensitive to outlying or unusually small variance estimates. To improve the stability of the substrate-level inference, we therefore used the variance adaptive shrinkage (VASH) empirical Bayes method [38]. VASH borrows information across all retained substrates to estimate the overall distribution of variances, while still allowing each substrate to retain its own variance estimate.

For substrate $j$, let $\hat{s}_j^2$ denote the estimated variance associated with its effect estimate, $s_j^2$ the corresponding unknown true variance, and $d_j$ the residual degrees of freedom. VASH uses the standard sampling model

$$\hat{s}_j^2 \mid s_j^2 \sim s_j^2 \frac{\chi^2_{d_j}}{d_j}.$$

The true variances across substrates are assigned a flexible unimodal empirical Bayes prior, represented as a mixture of inverse-gamma distributions constrained to share a common mode. The prior mode and mixture proportions are estimated from the observed variance estimates across all retained substrates by maximizing the marginal likelihood, so the degree of shrinkage is learned from the data rather than fixed in advance.

For each substrate, the observed variance estimate is combined with the fitted cross-substrate prior to obtain a posterior mixture of inverse-gamma distributions. Following Lu and Stephens [38], we used the inverse of the posterior mean precision,

$$\tilde{s}_j^2 = \left\{ \mathrm{E}\left( s_j^{-2} \mid \hat{s}_j^2 \right) \right\}^{-1},$$

as the moderated variance estimate. Substrate-level two-sided $p$-values were then calculated using the corresponding VASH moderated $t$-test. This empirical Bayes moderation reduces the influence of unstable raw variance estimates, particularly spuriously small estimates that can arise in small-sample settings, while preserving substrate-specific heterogeneity.

The resulting substrate-level $p$-values and corresponding network annotations are provided in the Supplementary Table S3.

### Supplementary Material 3: SCZ Substrate Distributions and Sensitivity Analysis

#### Phosphosite list

The list of phosphosites was generated from phosphopeptide data, using in-house R scripts adapted from Wozniak et al. [40]. Phosphosites were only retained with a localization score of > 0.75 and mapped to their corresponding position with the protein using human reference proteome UP000005640 [34], as a reference. To map phosphosites to kinase/substrate interactions, KSEA [39] was utilized, with databases (Phosphosite [36] PLUS, NetworKIN [37]), providing a final list of kinase/phosphosites.

#### Differential phosphorylation

Supplementary Fig. S1 shows the schizophrenia case-control differential phosphorylation distributions for phosphosites downstream of MAPK3, CSNK2A1, and CSNK2A2, as discussed in the main text.

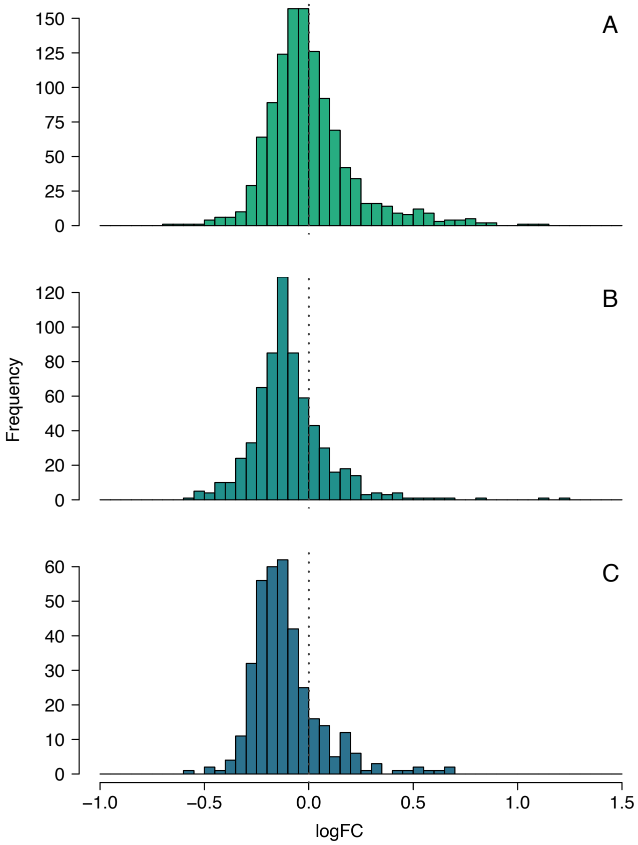


**Fig. S1.** Distribution of differential phosphorylation among predicted downstream phosphosites for MAPK3, CSNK2A1, and CSNK2A2 in the schizophrenia dataset. Panel a shows MAPK3-associated phosphosites, and panels b–c show phosphosites associated with the CK2 catalytic subunits CSNK2A1 and CSNK2A2. The MAPK3 distribution shows a long right tail, whereas the CK2-associated phosphosites tend to be shifted toward lower phosphorylation in schizophrenia cases.

#### Sensitivity analysis for LIKA

We assessed whether LIKA kinase rankings are sensitive to the first-stage substrate-level FDR threshold. In the LIKA workflow, empirical Bayes phosphosite $p$-values are thresholded using the Benjamini–Hochberg procedure to define the significant phosphosite set used for kinase-level inference. A robust ranking should therefore remain stable across reasonable FDR choices.

For the INKA and SCZ datasets, we repeated LIKA using first-stage FDR levels in $\{0.01,\ 0.025,\ 0.05,\ 0.10,\ 0.20\}$.

At each threshold, we recomputed the significant phosphosite set and used the largest raw substrate $p$-value among rejected phosphosites as the fixed null probability in the LIKA kinase-level test. Kinases were ranked by capped LIKA $p$-value, defined as $\max(p, 0.05/m)$, where $m$ is the number of kinases with valid LIKA $p$-values; ties were broken by the number of efficient substrates. We compared rankings across FDR levels using the top-10 overlap ratio, $\frac{|T_a \cap T_b|}{10}$, where $T_a$ and $T_b$ denote the top-10 kinase sets from two FDR thresholds.

Supplementary Fig. S2 shows the resulting top-10 overlap ratios across FDR thresholds.

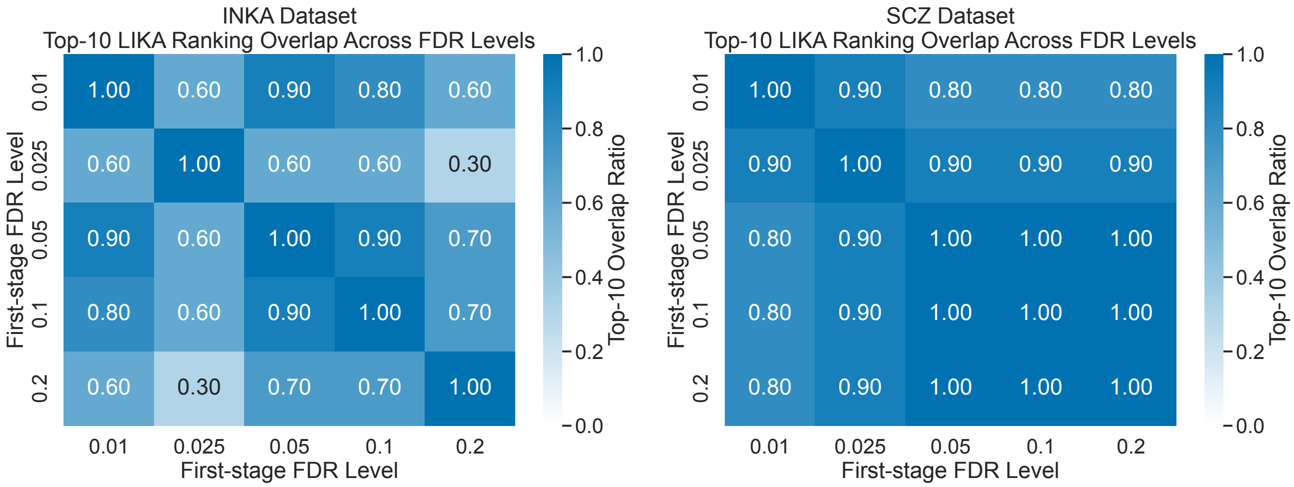


**Fig. S2.** Sensitivity of LIKA top-10 kinase rankings to the first-stage FDR threshold. Each heatmap entry shows the top-10 overlap ratio, $|T_a \cap T_b|/10$, between two FDR settings. The left panel shows the INKA dataset and the right panel shows the SCZ dataset; darker blue indicates stronger agreement. SCZ rankings were highly stable, with all pairwise overlaps at least 0.80 and complete overlap among FDR 0.05, 0.10, and 0.20. INKA rankings were moderately stable but more threshold-dependent, with off-diagonal overlaps ranging from 0.30 to 0.90.

Overall, SCZ top-ranked LIKA kinases were largely insensitive to the first-stage FDR threshold. INKA showed greater threshold dependence, particularly between stringent and relaxed thresholds, but the default FDR 0.05 ranking remained close to neighboring settings, with 90% top-10 overlap against both FDR 0.01 and FDR 0.10.
